# Single-cell RNA-seq of the Developing Cardiac Outflow Tract Reveals Convergent Development of the Vascular Smooth Muscle Cells at the Base of the Great Arteries

**DOI:** 10.1101/469346

**Authors:** Xuanyu Liu, Wen Chen, Wenke Li, James R. Priest, Jikui Wang, Zhou Zhou

**Author notes:** **Subject Terms**: Developmental Biology, Vascular Biology, Gene Expression and Regulation, Congenital Heart Disease. **Address correspondence to:** Dr. Jikui Wang, School of Basic Medical Sciences, Xinxiang Medical University, No. 601, Jinsui Road, 453003 Xinxiang, China; Dr. Zhou Zhou, Fuwai Hospital, Chinese Academy of Medical Sciences, No. 167 Beilishi Street, Xicheng District, 100037 Beijing, China.

## Abstract

**Rationale:** Cardiac outflow tract (OFT) is a major hotspot for congenital heart diseases (CHDs). A thorough understanding of the cellular diversity, transitions and regulatory networks of normal OFT development is essential to decipher the etiology of OFT malformations.

**Objective:** We sought to explore the cellular diversity and transitions between cell lineages during OFT development.

**Methods and Results:** We performed single-cell transcriptomic sequencing of 55,611 mouse OFT cells from three developmental stages that generally correspond to the early, middle and late stages of OFT remodeling and septation. We identified 17 cell clusters that could be assigned to six cell lineages. Among these lineages, the macrophage and VSMC lineages of the developing OFT have seldom been previously described. Known cellular transitions, such as endothelial to mesenchymal transition, have been recapitulated. In particular, we identified convergent development of the VSMC lineage, where intermediate cell subpopulations were found to be involved in either myocardial to VSMC trans-differentiation or mesenchymal to VSMC transition. Through single-molecule *in situ* hybridization, we observed that cells expressing the myocardial marker *Myh7* co-expressed the VSMC marker gene *Cxcl12* in OFT walls, thus confirming the existence of myocardial to VSMC trans-differentiation. Moreover, we found that the *Penk*^+^ cluster c8, a relatively small mesenchymal subpopulation that was undergoing mesenchymal to VSMC transition, was associated with the fusion of OFT cushions. We also uncovered the expression dynamics and critical transcriptional regulators potentially governing cell state transitions. Finally, we developed web-based interactive interfaces to facilitate further data exploration.

**Conclusions:** We provide a single-cell reference map of cell states for normal OFT development, which will be a valuable resource for the CHD community. Our data support the existence of myocardial to VSMC trans-differentiation and convergent development of the VSMC lineage at the base of the great arteries.

## INTRODUCTION

Congenital heart disease (CHD) is the most common form of human birth defects (∼1% of live births) and represents the leading cause of mortality from birth defects worldwide^1^. Approximately 30% of CHDs involve abnormalities in cardiac outflow tract (OFT) development, thus constituting a large class of CHDs, namely OFT malformations, such as persistent truncus arteriosus (PTA), double outlet right ventricle (DORV), transposition of the great arteries (TGA) and aortopulmonary-window (APW)^2, 3^. It is therefore acknowledged that OFT is a major hotspot for human CHDs^4^. OFT malformations require surgical repair once diagnosed and usually have a poor prognosis^5^. However, the etiology for the majority of this severe class of CHDs remains unknown.

Cardiac OFT is a transient conduit during embryogenesis at the arterial pole of the heart, connecting the aortic sac with embryonic ventricles, which undergoes rotation and septation (a.k.a., OFT remodeling) to give rise to the base of the pulmonary trunk and ascending aorta; thus, this process is critical for the establishment of separate systemic and pulmonary circulations^6^. The high incidence of OFT malformations may be explained by the complexity of OFT development, which requires intricate interplay and transitions among diverse cell populations, including cardiac cells and migrating extra-cardiac cells, making it particularly susceptible to genetic or environmental perturbations. A thorough understanding of the cellular diversity, cellular transitions and regulatory networks of normal OFT development is essential to decipher the etiology of OFT malformations.

Multiple disparate cell types have been implicated in OFT development through tightly coordinated processes such as migration, differentiation and transition. Initially, the OFT wall basically consists of a solitary tube of myocardium derived from the second heart field (SHF)^7^. At the initiation of remodeling, the interstitial space between the myocardium and endothelium is filled with extracellular matrix (“cardiac jelly”) secreted by mesenchymal cells that form OFT cushions at the proximal and distal regions^8^. Cardiac neural crest cells (CNCCs) migrate into the cardiac OFT, where they first join the mesenchyme of the distal and then the proximal cushions, playing an essential role in the fusion of the distal cushions to form a smooth muscle septum, i.e., aorticopulmonary septum, which divides the aorta and pulmonary trunk^9, 10^. In addition to CNCCs, distal OFT cushions are colonized by cells derived from the SHF, which eventually give rise to smooth muscle walls of the base of the great arteries^11^. In contrast, proximal OFT cushions are mainly populated by the mesenchymal progenies of OFT endothelial cells that undergo endothelial to mesenchymal transition (EndoMT)^12^. Besides, the OFT remodeling is accompanied by other biological processes, for example, the maturation of the smooth muscle walls, since the OFT wall changes from a myocardial to an arterial phenotype with development^13^. Given the complexity of OFT development, we expect a heterogeneous cellular composition represented by multiple subpopulations of the same cell type and extensive cellular transitions occurring between different cell types. However, the cell type and cell states of the cardiac OFT during development have not yet been systematically dissected.

Recent technical advances have enabled the transcriptomes of tens of thousands of cells to be assayed at single-cell resolution in a single experiment^14^. Single-cell RNA-seq has shown itself to be a powerful tool to provide insights into the processes underlying developmental, physiological and disease systems^15^. Single-cell RNA-seq enables the dissection of cellular heterogeneity in an unbiased manner with no need for any prior knowledge of the cell population^16^. Unsupervised clustering of cells based on genome-wide expression profiles enables the identification of novel cell types or subpopulations, as well as gene signatures for all cell types. Beyond cellular heterogeneity dissection, single-cell RNA-seq data empower systematic interrogations of the developmental trajectory of cell lineages in tissue systems and the regulatory networks underlying the cell state transition processes^17^. Although traditional gene knockdown studies have uncovered regulators during OFT development^10^, how genes are regulated under a normal developmental context remains unclear. Single-cell RNA-seq has been applied to study the cellular diversity of embryonic^18, 19^ or adult heart^20, 21^ at the whole organ level; however, few or a limited number of cardiac OFT cells have been sampled in these studies.

Here, we performed single-cell transcriptomic sequencing of 55,611 mouse OFT cells from three successive developmental stages corresponding to the early, middle and late stages of OFT remodeling and septation. We sought to unbiasedly and systematically dissect the cell types and states during OFT development. We explored the cell lineage relationships and cellular state transitions during OFT development, as well as the critical transcription regulators underlying the transitions. We identified convergent development of the vascular smooth muscle cells (VSMCs) at the base of the great arteries, where intermediate cell subpopulations were found to be involved in either myocardial to VSMC trans-differentiation or mesenchymal to VSMC transition. Our study provided a single-cell reference map of cell transcriptomic states for OFT normal development.

## METHODS

See **Online Methods**.

The sequencing read data have been deposited in Genome Sequence Archive (GSA; http://gsa.big.ac.cn/) and are accessible through accession number CRA001120.

## RESULTS

### Single-cell transcriptomic sequencing and unbiased clustering of developing OFT cells

To obtain a map of the cellulome for the developing OFT during remodeling and septation, we isolated and sequenced a total of 64,605 cells from three successive developmental stages, namely, ps47 (47 pairs of somites), ps49 and ps51, which generally correspond to the early (initiation), middle and late (almost completion) stages of septation, respectively (Figure 1A). Sequencing quality metrics were similar across samples, reflecting little technical variation among samples (Online Table I). After stringent quality filtering and discarding a small number (297) of red blood cells, we obtained a high-quality transcriptomic dataset for 55,611 cells.

**Figure 1.**
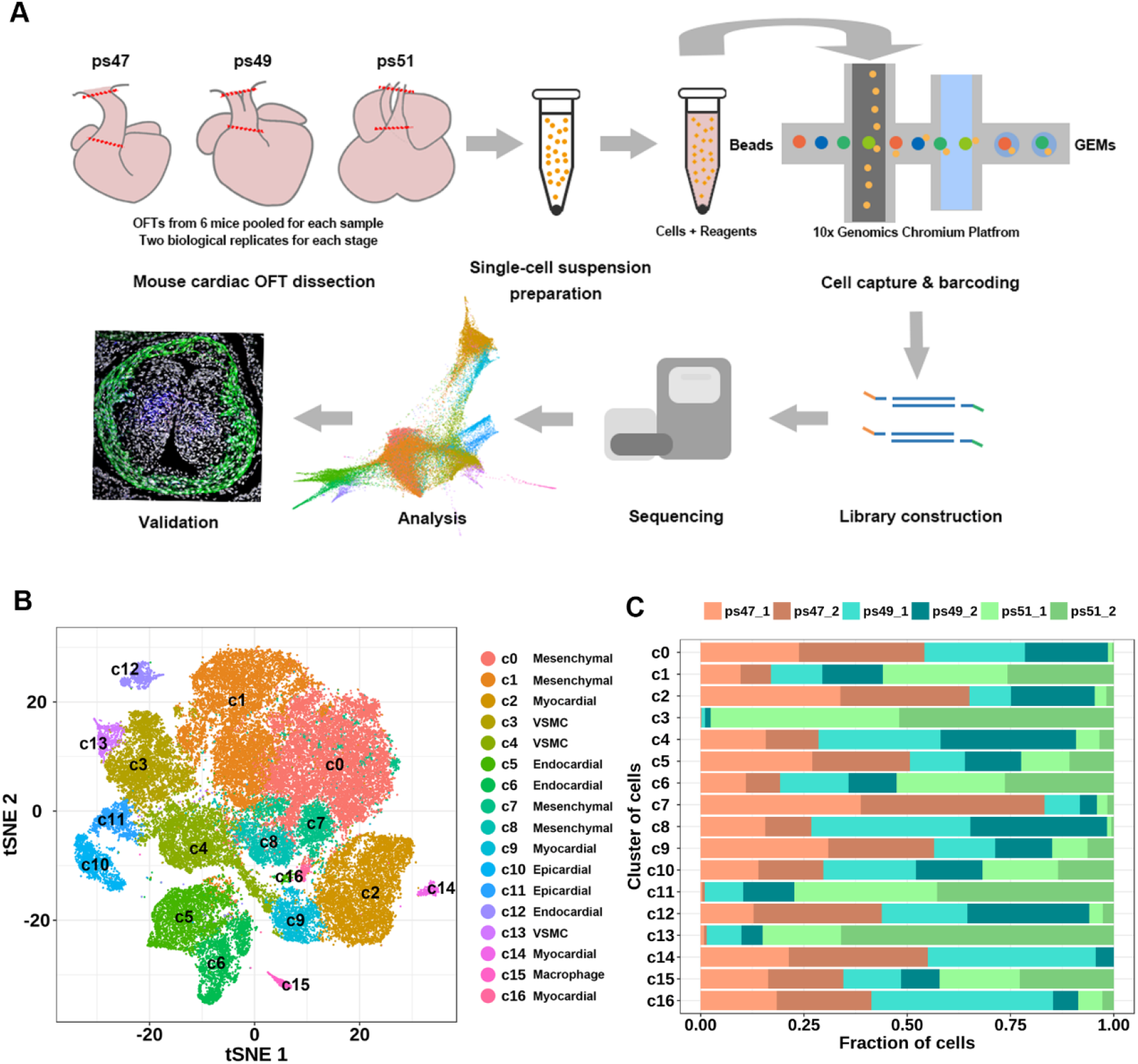
Single-cell transcriptomic sequencing and unbiased clustering of cells during OFT development. **(A)** Overview of the experimental procedure. Single-cell suspensions from three successive stages of mouse OFT development were captured and sequenced separately. Two biological replicates were prepared for each stage. ps47, ps49 and ps51 denote 47, 49 and 51 pairs of somites, respectively. Dissection boundaries are indicated by the red dotted lines on the schematic plots of embryonic hearts. **(B)** Unsupervised clustering of all cells reveals 17 cell clusters projected on a two-dimensional tSNE map. **(C)** Fraction of cells derived from each sample for each cluster. All samples are normalized to the same number of cells (4,270).

Based on the single-cell transcriptional profiles, unsupervised clustering identified 17 cell clusters at the chosen resolution (Figure 1B) that represent distinct cell types or cell subpopulations. Next, we compared the relative proportions of cells from different samples in each cluster (Figure 1C). Importantly, there was no significant difference in cell fraction between the two biological replicates for each stage, reflecting the validity of cell clustering (Wilcoxon signed rank test P-value = 0.25). Moreover, all the clusters contained cells from the three stages except c14, a small cluster (342 cells), suggesting that the samples for each stage covered all common cell states throughout development. However, the relative proportions varied greatly between stages, as reflected by the observation that the cell fractions for some clusters, e.g., c11, remarkably changed during development (Figure 1C).

### Cellular diversity and heterogeneity during OFT development identified by single-cell transcriptomic analysis

To define the identity of each cell cluster, we performed differential expression analysis between each cluster and all others (Online Table II), and assigned a specific cell type to each cluster based on the established lineage-specific marker genes (Figure 2A). Cluster c15 represented a small group of macrophages that resided in the developing OFT as the cells in this cluster specifically expressed *Fcgr1* and *Adgre*^21, 22^. Clusters c10 and c11 constituted the epicardial lineage as they specifically expressed *Upk3b* and *Upk1b*^19, 23^. Clusters c5, c6 and c12 highly expressed *Ecscr* and *Cdh5*^19, 24^; thus, they belonged to the endocardial lineage. Clusters c2, c9, c14 and c16 highly expressed myocardial marker genes, such as *Myh7* and *Myl4*^19, 25^. The mesenchymal lineage comprised four closely aligned clusters, namely, c0, c1, c7 and c8, which highly expressed mesenchymal marker genes, such as *Postn* and *Cthrc1*^19^. The VSMC lineage included c3, c4 and c13, which specifically expressed *Rgs5*, a gene that is abundantly expressed in arterial smooth muscle cells^26, 27^, and *Cxcl12*, a chemokine encoding gene that is highly expressed in the walls of the aorta and pulmonary trunk of the embryonic heart (E12.5)^28^. We also assessed the expression intensity distribution of contractile markers for smooth muscle cells, including *Acta2*, *Tagln*, *Cnn1* and *Myl9*^25, 29^. We observed the expression of these markers in our VSMC lineage clusters. However, the specificity of these markers was not ideal since they were also highly expressed in myocardial and mesenchymal lineages (Online Figure IA), which is in line with previous knowledge^29^. Moreover, only a small group of cells expressed *Myh11*, the most specific contractile marker for mature VSMCs^29^, and clustered on the edge of c3, which may represent a group of relatively mature VSMCs. By contrast, the expression of contractile markers for embryonic myocardial cells^25^, including *Myh7*, *Tnnc1*, *Tnnt2*, *Myl2* and *Myl4*, were relatively specific in myocardial lineage clusters (Online Figure IB). Ultimately, we identified clusters of VSMCs at the base of the great arteries, most of which may exhibit an immature, synthetic phenotype at this developmental stage.

**Figure 2.**
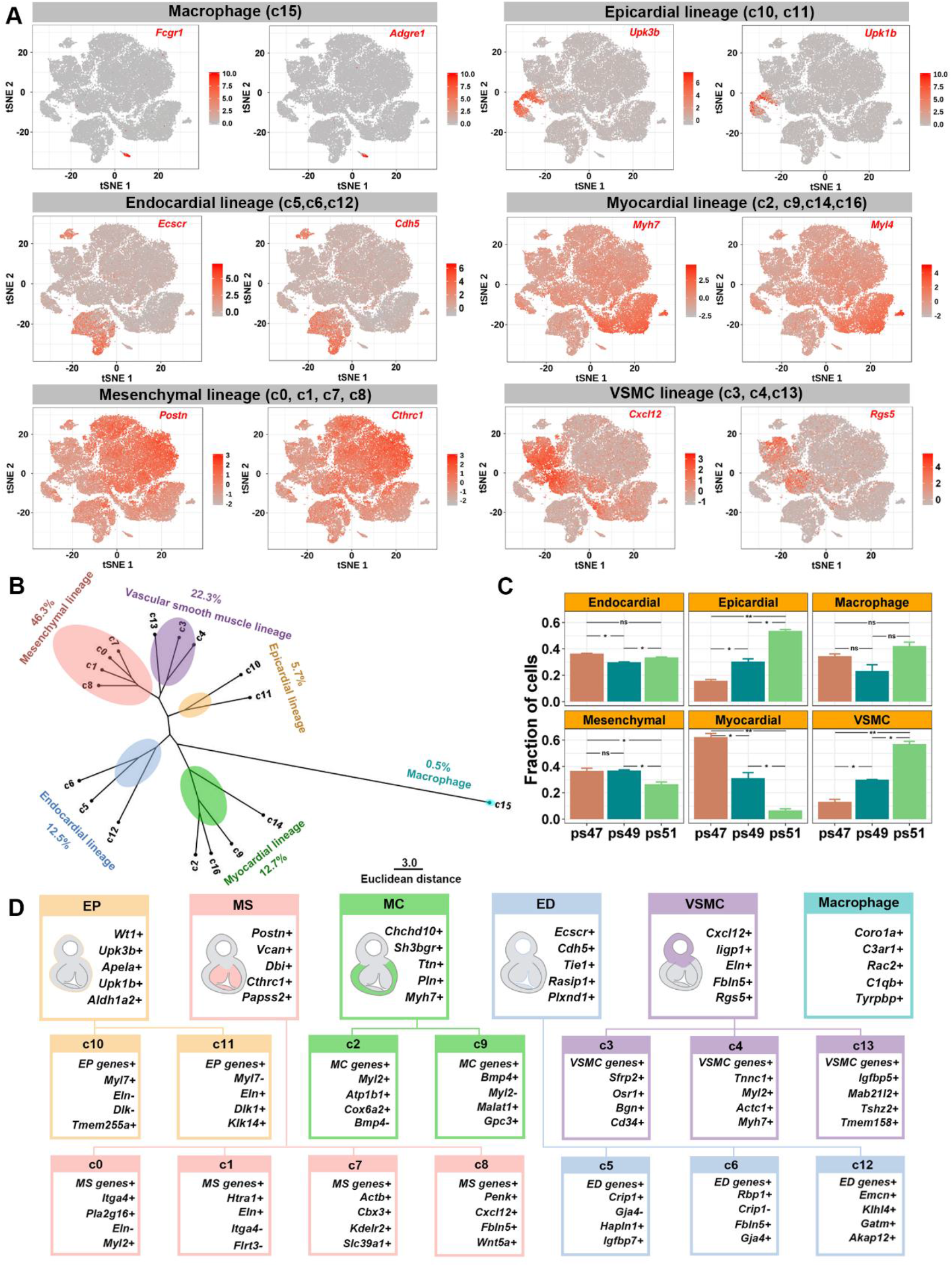
Cellular diversity and gene signatures of the developing OFT identified by single-cell transcriptomic analysis. **(A)** Cell lineages recognized by known cell-type specific marker genes. Each cell is colored according to the scaled expression of the indicated marker gene. **(B)** Relatedness of clusters revealed by hierarchical clustering. This analysis is based on the average expression of the 1,381 HVGs in each cluster. **(C)** Cell fractions of each stage for each cell lineage. The average cell fraction of the two biological replicates and the standard error are shown on the bar plot. ‘ns’: not significant; *: Student's t-Test P-value < 0.05; **: Student's t-Test P-value < 0.01. In B and C, all samples are normalized to the same number of cells (4,270). **(D)** Gene signature for each cell lineage or cluster. These genes were identified and selected by differential expression analysis and random forest classification. The schematic plot represents a cross section of OFT in a specific position where the aorta wall shows an artery phenotype while the wall of the pulmonary artery still possesses a myocardial phenotype. The two small myocardial clusters c14 and c16 were not incorporated in this analysis. EP: epicardial; MS: mesenchymal; MC: myocardial; ED: endocardial; VSMC: vascular smooth muscle cell

The six cell lineages identified by established marker genes were further confirmed by hierarchical clustering analysis, which showed that the clusters assigned to the same lineage were grouped together and closely aligned on the tree (Figure 2B). Once cell identity was assigned, we explored the relative proportions of each cell lineage during development (Figure 2B & 2C). The mesenchymal lineage in OFT cushions constituted the most abundant cell type (46.3%), and the relative proportion significantly decreased at the late stage (Student's t-Test P-value < 0.05), suggesting that an active cellular transition occurred at the late stage. The macrophages accounted for only 0.5% of all cells, and the relative proportion did not significantly change during development. Strikingly, the myocardial lineage diminished over time, while the VSMC lineage expanded during development, in accordance with the myocardial to arterial phenotypic change. The epicardial lineage also significantly expanded with development, consistent with a previous report^19^.

### Machine learning-based selection of molecular signatures for cell lineages and clusters during OFT development

To select molecular signatures that define the identified cell lineages and clusters, we adopted a machine learning-based strategy (Online Figure II). All the random forest models we trained achieved a good classification performance (AUC range: 0.94–1; Online Figure III & Online Figure IV). The top genes that contributed the most to the models were selected as the molecular signatures for each of the six cell lineages or each subpopulation/cluster (Figure 2D). For cell lineages, most selected genes have been reported to be specifically expressed in a cell type. For example, *Wt1* in epicardium^30^, *Vcan* in cushion mesenchyme^31^, *Myh7* in myocardium^32^, *Rasip1* in endocardium^33^ and *Fbln5* in VSMCs^34^. However, some genes have seldom been previously described to be expressed specifically in a cell lineage of the developing OFT, e.g., *Aldh1a2* in epicardium and *Papss2* in mesenchyme. For clusters of each lineage, some selected markers have been reported to be cell type-specific in the embryonic heart (E10.5)^19^; however, they were found to be expressed in a cluster-specific manner. For example, *Tmem255a* was reported to be the epicardium marker of the E10.5 heart^19^; however, in the present study, it was specifically expressed in only one of the two subpopulations of the epicardial cells, namely, c10 (Online Figure IV).

### Convergent development of the VSMCs at the base of the great arteries inferred from a KNN graph

To infer the relationships of cell lineages, we visualized the single-cell dataset using a force-directed layout of a k-nearest-neighbor (KNN) graph (Figure 3A & 3B), which has been proven to perform better than t-distributed stochastic neighbor embedding (tSNE) for visualizing complex and continuous gene expression topologies of cell populations^35^. Cell clusters of the same lineage were closely aligned in the KNN graph, and known relationships among lineages have been well recapitulated. For example, the EndoMT process was reflected by an abundance of potentially intermediate, transitioning cells connecting the endocardial lineage (c5) and mesenchymal lineage (c1), while these cell clusters were separated in the tSNE plot shown in Figure 1B. The dynamic changes in cell states during development could be well visualized when the cells were displayed by developmental stage (Figure 3C). For example, we observed the rapidly diminished myocardium and the expanded epicardium.

**Figure 3.**
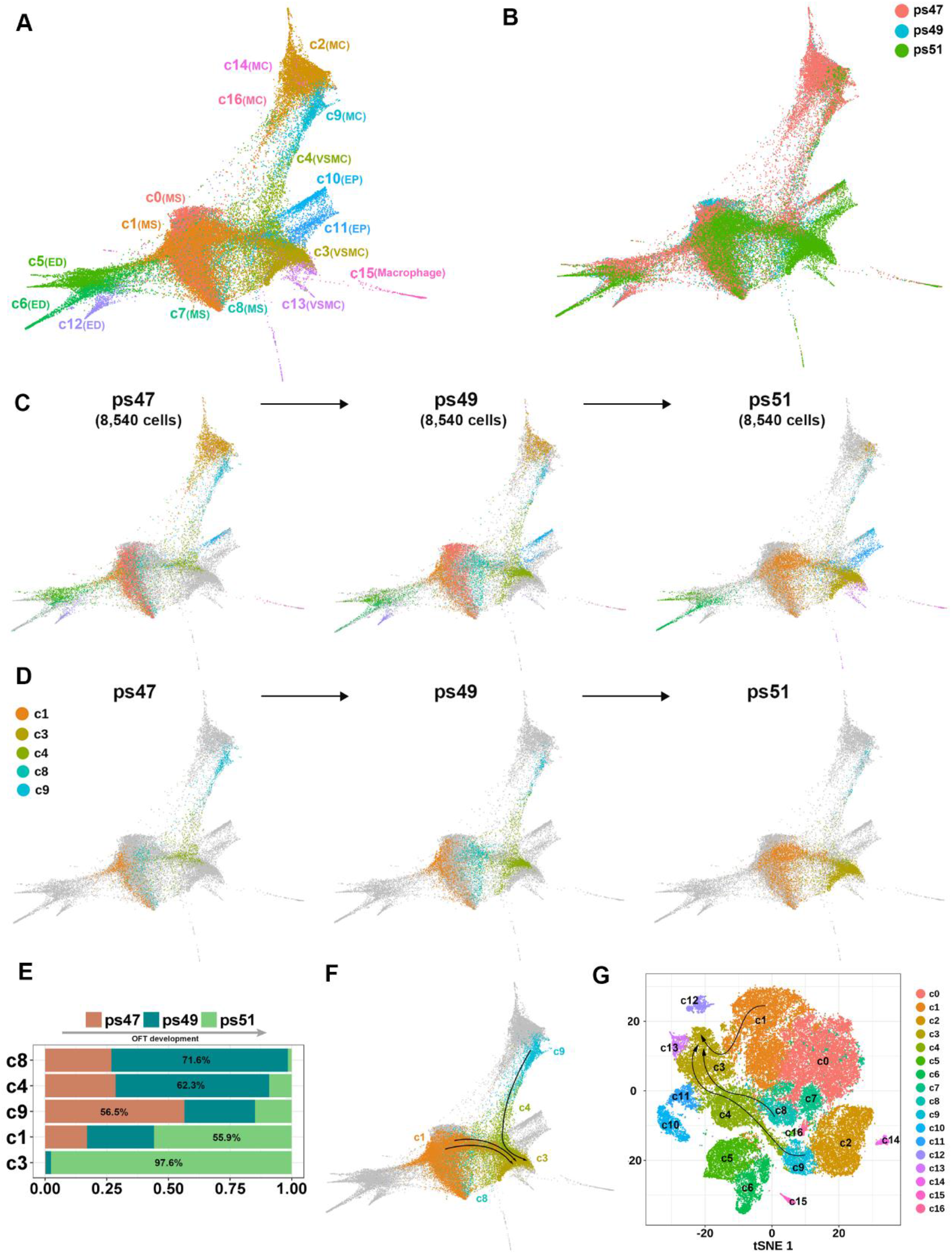
Convergent development of the VSMCs at the base of the great arteries. **(A)** Force-directed layout of a KNN graph showing a continuous expression topology of the OFT cellulome during development. Each dot denotes a cell colored by cell cluster. **(B)** KNN graph colored by development stage. **(C)** Dynamic changes of cell states over time. Cells colored in gray denote those from other stages. All samples are normalized to the same number of cells (4,270). **(D)** Dynamic changes of cell clusters directly involved in VSMC development. **(E)** Relative proportions of cell clusters directly involved in VSMC development. **(F)** Inferred developmental paths of the smooth muscle cells displayed on the KNN graph. **(G)** Inferred developmental paths of the smooth muscle cells displayed on the tSNE plot.

In particular, we noticed five cell clusters, including c1, c3, c4, c8 and c9, which may be directly related to the development of VSMCs (Figure 3D & 3E). VSMC cluster c4 expanded during the early and middle stage and was almost replaced by VSMC cluster c3 at the late stage. In the KNN plot (Figure 3D), c4 became closer to c3 over time. Therefore, we speculated that c4 may represent an intermediate state and c3 a more mature state of VSMCs. Intriguingly, myocardial subpopulation c9 and VSMC progenitor population c4 were densely connected, and a considerable number of cells were in between, which may represent intermediate, transitioning cell states. Therefore, this finding could imply that myocardial to VSMC trans-differentiation may occur during OFT development. Moreover, a relatively small (1,924 cells) mesenchymal cluster, c8, was closely aligned with c4 and became closer to c4 over time, implying that c8 may represent a special mesenchymal subpopulation actively involved in the mesenchymal to VSMC transition at a relatively early stage. Additionally, c1, a relatively large (10,407 cells) mesenchymal subpopulation may also be involved in the mesenchymal to VSMC transition, particularly at a relatively late stage, as reflected by an abundance of potentially intermediate cells connecting.c1 and c3, mainly at stage ps51 (Figure 3D).

Altogether, our data suggest convergent development of the VSMCs at the base of the great arteries, where intermediate cell subpopulations were found to be involved in either mesenchymal to VSMC transition or myocardial to VSMC trans-differentiation. The inferred development paths of VSMCs are summarized in Figure 3F. Since a considerable number of intermediate cells were captured in our dataset, these biological processes could even be directly inferred from the tSNE plot (Figure 3G).

### Characteristics of gene expression profiles for intermediate cell subpopulations involved in the myocardial to VSMC trans-differentiation

We next sought to confirm myocardial to VSMC trans-differentiation by examining the gene expression profiles of the intermediate cell subpopulations that we identified above, c9 and c4. Cell cluster c9 and the largest myocardial cluster c2 displayed distinct expression profiles (Figure 4A). Compared with c2, c9 showed significant up-regulation of VSMC marker genes, including contractile VSMC markers^29^ (e.g., *Acta2*, *Tagln*, *Cald1* and *Myl9*) and synthetic VSMC markers^36^ (e.g., *Eln, Col1a2 and Cxcl12*; Figure 4A, B). Myocardium-specific genes, *Tnnc1* and *Tnnt2*^25^, were expressed at significantly higher levels in c9 than in c2, reflecting the myocardial identity of c9. However, some myocardial markers, such as *Myl2*, were significantly down-regulated in c9. Moreover, the ratio of *Myh6* to *Myh7*, an index reflecting the degree of maturation and functionality of cardiomyocytes^37^, was significantly lower in c9 (Wilcoxon rank sum test P-value < 2.2e-16; Figure 4C), suggesting that c9 cells may exhibit a less “mature” phenotype and be undergoing phenotypic changes. The genes up-regulated in c9 versus c2 (Online Table III) were enriched for pathways such as smooth muscle contraction, elastic fiber formation and artery morphogenesis (Figure 4D).

**Figure 4.**
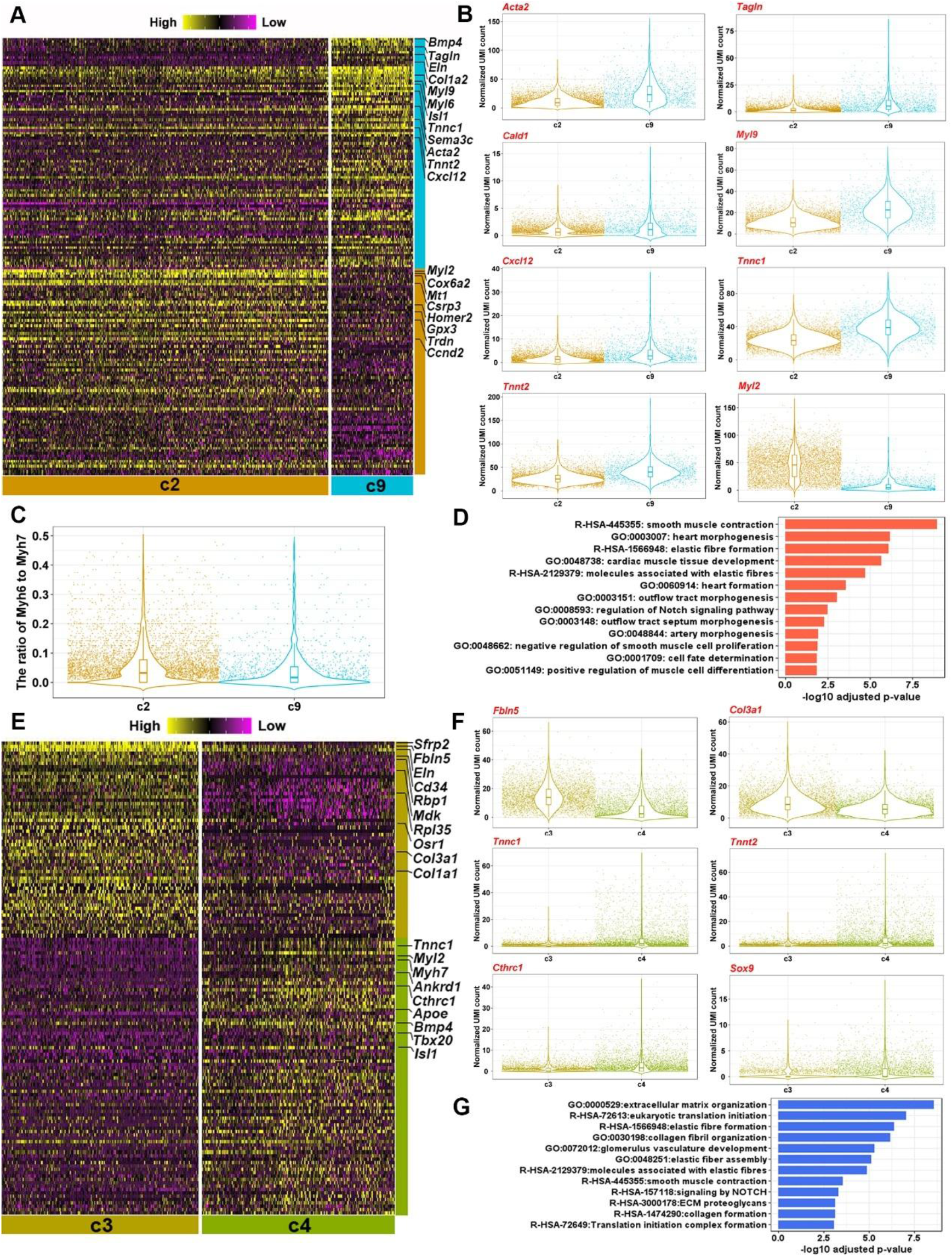
Characteristics of gene expression profiles for the intermediate cell subpopulations during myocardial to VSMC trans-differentiation. **(A)** Heatmap showing the DEGs between myocardial cluster c2 and c9. **(B)** Significant expression differences of some key marker genes between c2 and c9. Each dot denotes a cell. **(C)** The ratio of Myh6 to Myh7 in c2 and c9. **(D)** Functional enrichment of genes significantly up-regulated in c9. **(E)** Heatmap showing the DEGs between the VSMC cluster c3 and the intermediate VSMC population c4. **(F)** Significant expression differences of some key marker genes between c3 and c4. **(G)** Functional enrichment of genes significantly up-regulated in c3.

We also observed distinct expression profiles between the two VSMC clusters, c3 and c4 (Figure 4E, Online Table IV). Genes related to the development and maturation of VSMCs, such as *Fbln5*, *Eln*, *Col3a1* and *Col1a1*, were significantly up-regulated in c3 versus c4 (Figure 4E & 4F). The genes up-regulated in c3 were mainly enriched for ECM organization (Figure 4G). These results support our inference that c3 represents a more mature state of VSMCs than c4 does. Compared with c3, c4 exhibited significantly higher expression of both myocardial markers (e.g., *Tnnc1* and *Tnnt2*^25^) and mesenchymal markers (e.g. *Cthrc1* and *Sox9*^19^), reflecting its myocardial and mesenchymal heritage.

Altogether, the expression profile analysis supports our inference that c9 and c4 cells are in an intermediate state along the trajectory of myocardium to VSMC trans-differentiation.

### Pseudo-temporal ordering and gene regulatory network analysis uncover critical transcriptional regulators potentially governing cell state transitions during OFT development

To elucidate gene expression dynamics, especially the transcriptional regulators governing the convergent development of VSMCs, we reconstructed the development trajectories for the different paths (Figure 3G) through pseudo-temporal ordering of individual cells using CellRouter (See Methods). For the myocardial to VSMC trans-differentiation (c9-c4-c3), we identified genes that were significantly correlated with the trajectory (Online Table V) and observed the loss of myocardial marker expression and gain of VSMC marker expression during the progression of trans-differentiation (Figure 5A). The Notch signaling pathway positively regulates the specification, differentiation, and maturation of VSMCs^38^. Strikingly, we noted that the expression of genes in the Notch signaling pathway, including receptor (*Notch1*), ligand (*Jag1*) and downstream targets (*Hey1*, *Hey2*, *Heyl* and *Pdgfrb*), was positively correlated with the trajectory (Figure 5B). Furthermore, by gene regulatory network (GRN) analysis, we obtained a set of key regulators that were activated sequentially along the trajectory and potentially drove the process (Figure 5C). We identified *Heyl*, encoding a known downstream TF of Notch signaling, at the top of the up-regulated regulators, consistent with its known positive role in VSMC development^38^. *Tbx20*, encoding a TF known as a transcriptional repressor in the developing heart^39^, was at the top of the down-regulated regulators, implying its role in repressing VSMC lineage-specifying genes. Regulators that were activated early in the trajectory, such as *Plagl1* and *Naca,* may play potential roles in lineage commitment.

**Figure 5.**
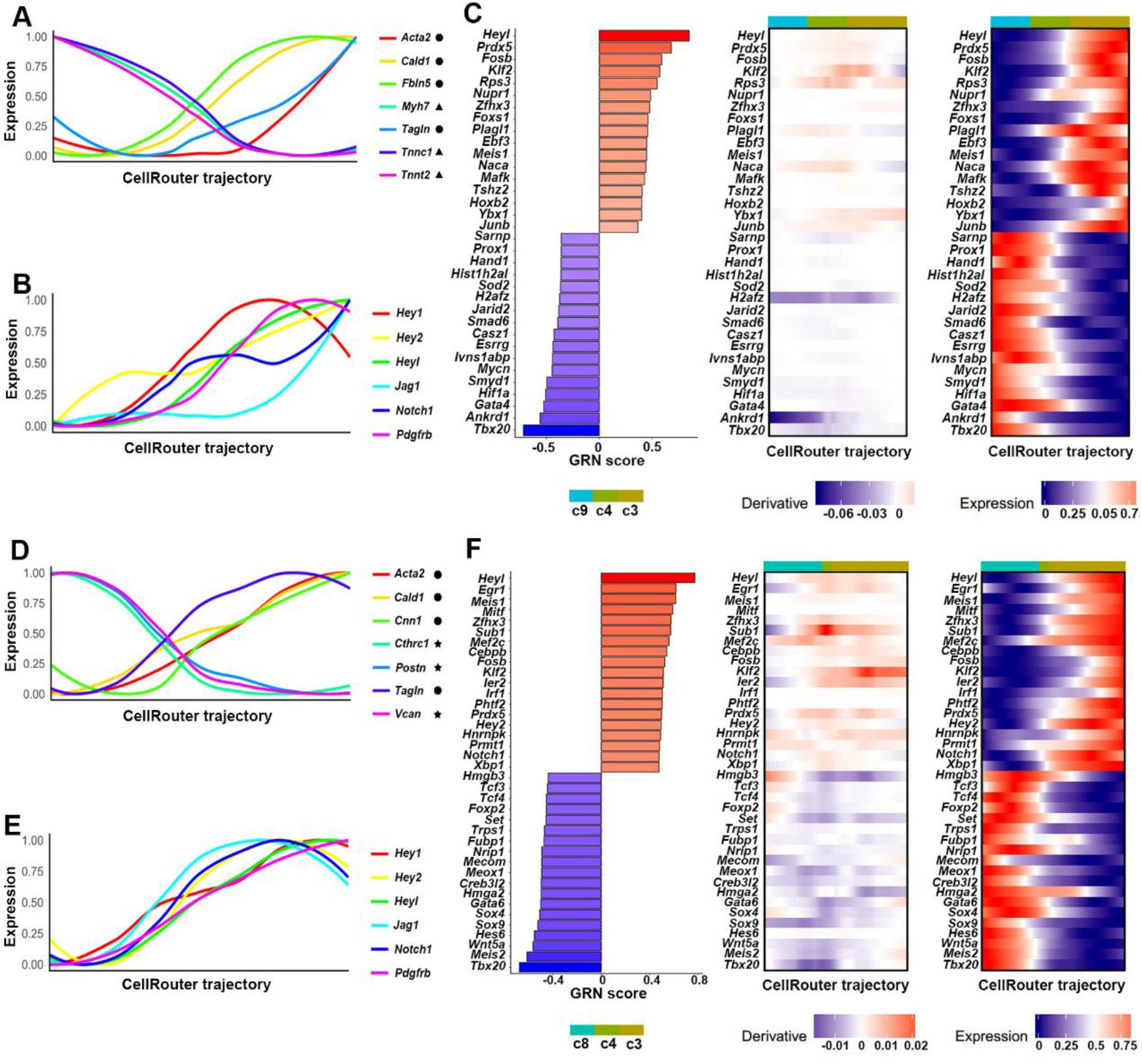
Pseudo-temporal ordering and GRN analysis uncover critical transcriptional regulators potentially governing cell state transitions during the development of VSMCs. **(A)** Loss of myocardial marker expression and gain of VSMC marker expression during myocardial to VSMC trans-differentiation. **(B)** Expression of genes in the Notch signaling is positively correlated with the trajectory of myocardial to VSMC trans-differentiation. **(C)** Critical transcriptional regulators potentially involved in myocardial to VSMC trans-differentiation. **(D)** Loss of mesenchymal marker expression and gain of VSMC marker expression during c8 mesenchymal to VSMC transition. **(E)** Expression of genes in the Notch signaling is positively correlated with the trajectory of c8 mesenchymal to VSMC transition. **(F)** Critical transcriptional regulators potentially involved in c8 mesenchymal to VSMC transition. ●: VSMC marker; ▲: myocardial marker; ★: mesenchymal marker.

For mesenchymal (the c8 *Penk^+^* subpopulation) to VSMC transition (c8-c4-c3), we identified genes that were significantly correlated with the trajectory (Online Table VI) and observed the loss of mesenchymal marker expression and gain of VSMC marker expression during the progression of the transition (Figure 5D). The expression of genes in the Notch signaling pathway was positively correlated with the trajectory (Figure 5E). Interestingly, we found *Heyl* and *Tbx20* also ranked at the top of the key regulators (Figure 5F). Consistent with our knowledge, the positive regulator *Mef2c*, a TF essential for VSMC development^40^, was activated relatively early in the reconstructed trajectory.

Similarly, for c1 mesenchymal to VSMC transition (c1_ps51_-c3), we observed the loss of mesenchymal marker expression and gain of VSMC marker expression during the progression of the transition (Online Figure VA, Online Table VII). The expression of genes in the Notch signaling pathway was positively correlated with the trajectory (Online Figure VB). By contrast, the top regulators were different from those in the c8 mesenchymal to VSMC transition (Online Figure VC).

Additionally, we reconstructed the trajectory and identified the gene expression dynamics for the EndoMT process between c5 endothelial cells and c1 mesenchymal cells (Online Table VIII, Online Figure VIA). We observed the loss of endocardial marker expression and gain of mesenchymal marker expression during the progression of the transition (Online Figure VIB). The expression of genes that have been implicated in EndoMT, particularly genes of the TGFβ signaling pathway^41^, were positively correlated with the trajectory (Online Figure VIC). Furthermore, we identified the critical transcriptional regulators potentially involved in the transition (Online Figure VID). Interestingly, *Klf2*, encoding a TF that may play a role in EMT during cardiac development^42^, was found to be ranked at the top of critical regulators. The predicted targets of *Klf2* were mainly enriched for epithelial to mesenchymal transition (Online Figure VIE), and included 16 genes that have been implicated in EMT (Gene Ontology term: epithelial to mesenchymal transition; Online Figure VIF).

### *Convergent development of the VSMCs at the base of the great arteries is confirmed by* single-molecule fluorescent *in situ* hybridization

To experimentally confirm myocardial to VSMC trans-differentiation, we performed single-molecule fluorescent *in situ* hybridization (smFISH) with probes for *Myh7* (myocardial marker), *Cxcl12* (VSMC marker) and *Bmp4* (the myocardial subpopulation c9 marker), which were selected based on our single-cell dataset (Figure 2D). Serial sections of the OFT at the middle stage ps49 from proximal to distal clearly showed myocardial to arterial phenotypic change in OFT walls (Figure 6A). Cells expressing high levels of *Myh7* (green) in OFT walls gradually changed into cells expressing *Cxcl12* (blue) over development. Remarkably, this change occurred faster on the aortic side than on the pulmonary arterial side. In addition to OFT walls, the expression of *Cxcl12* was specifically observed in a strip of cushion mesenchymal cells between the lumens of aorta and pulmonary artery, which developed into the aorticopulmonary septum, a smooth muscle structure that eventually forms the facing walls of the great arteries. These results illustrated that *Cxcl12* could serve as an early specific marker for the VSMC lineage of the OFT. In a single section, we could observe cells expressing myocardial marker *Myh7* co-expressed various levels of *Bmp4* and the *VSMC* marker gene *Cxcl12*, indicating a continuum of cell state transitions during myocardial to VSMC trans-differentiation (Figure 6B). The *Myh7*^+^*Cxcl12*^low^*Bmp4*^high^ cells that we observed (right panel of Figure 6B) may correspond to the c9 myocardial subpopulation.

**Figure 6.**
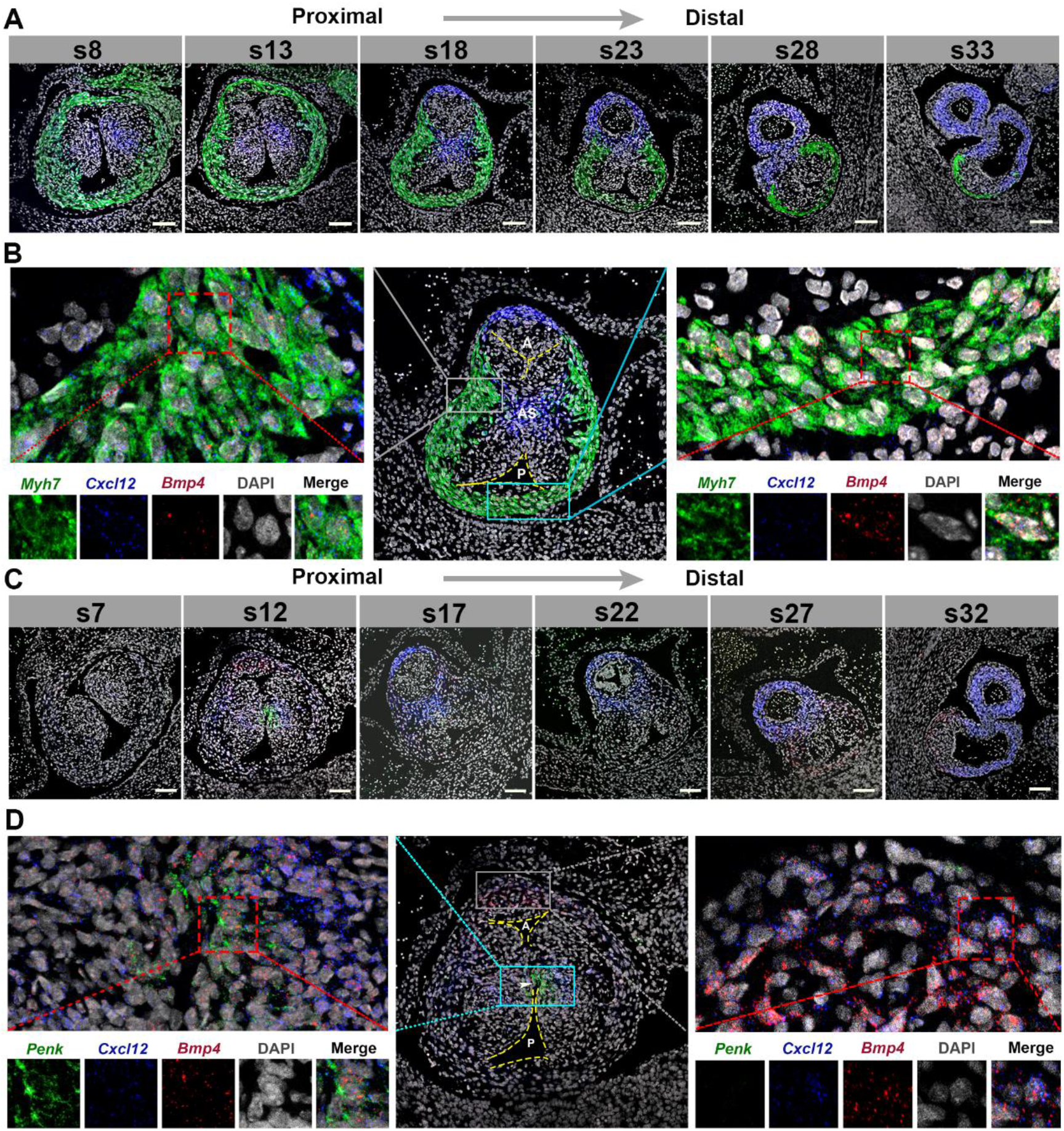
Convergent development of the VSMCs at the base of the great arteries is confirmed by smFISH. **(A)** Serial sections of the OFT at stage ps49 from proximal to distal showing myocardial to arterial phenotypic change in the OFT walls. Green: *Myh7*; Blue: *Cxcl12*; Red: *Bmp4*. Scale bar: 50 μm. The aorta is arranged on the upper side. **(B)** Myocardial to VSMC trans-differentiation is supported by the observation that cells expressing myocardial marker *Myh7* co-express various levels of *Bmp4* and VSMC marker gene *Cxcl12*. Middle panel: section S18; The yellow dotted line shows the border of the lumen of aorta (A) and pulmonary trunk (P); AS: aorticopulmonary septum. Left panel: *Myh7*^+^*Cxcl12*^high^*Bmp4*^low^ cells; Right panel: *Myh7*^+^*Cxcl12*^low^*Bmp4*^high^ cells. **(C)** Serial sections of the OFT at the stage ps49 from proximal to distal showing that the c8 *Penk*^+^ mesenchymal subpopulation is undergoing transition to VSMCs and is associated with the fusion of the OFT cushions. Green: *Penk*; Blue: *Cxcl12*; Red: *Bmp4*. Scale bar: 50 μm. **(D)** Co-expression of *Penk*, *Cxcl12* and *Bmp4*. The arrow indicates the location where the fusion is occurring. Middle panel: section S12; Left panel: *Penk*^+^*Cxcl12*^+^*Bmp4*^+^ cells; Right panel: *Penk^-^Cxcl12*^low^*Bmp4*^high^ cells.

Next, we tried to validate the existence of the c8 *Penk*^+^ mesenchymal subpopulation and confirm its role in mesenchymal to VSMC transition. Interestingly, serial sections of the OFT from proximal to distal showed that the expression of *Penk* could be observed only at the cushion mesenchyme where the fusion was occurring (the section S12, Figure 6C). We observed *Penk*^+^ mesenchymal cells co-expressed the VSMC marker *Cxcl12*^+^ and *Bmp4*^+^ (left panel of Figure 6D). These results suggest that the c8 *Penk*^+^ mesenchymal subpopulation is undergoing transition to VSMCs and may be associated with the fusion of the OFT cushions. Additionally, at section S12, the co-expression of *Bmp4* and *Cxcl12* could be observed at both the mesenchyme of the aorticopulmonary septum (left panel of Figure 6D) and the myocardial free wall of the aortic side (right panel of Figure 6D), which imply that independent of the developmental paths, *Bmp4* signaling may be associated with VSMC development.

### Web-based interfaces for further exploration of the single-cell data for the developing OFT

Our dataset constitutes a valuable resource for the scientific community to prioritize the candidate genes of OFT malformations based on expression and map the candidate genes to cell types/subpopulations. To facilitate further data exploration, we developed web-based interfaces for our dataset (http://singlecelloft.fwgenetics.org). These tools permit interactive examination of expression for any gene of interest, dynamic changes in cell states for each cluster in a 3D space, and potential intercellular communications among cell lineages for any ligand-receptor pair. Based on the expression of known ligand-receptor pairs, we observed extensive networks of potential intercellular communications among all cell lineages at each developmental stage (Online Figure VII). Interestingly, the network became significantly denser at the middle stage ps49 than at the other two stages, indicating increased intercellular communications at the middle stage.

## DISCUSSION

In the present study, we performed single-cell transcriptomic sequencing of 55,611 mouse OFT cells from three successive developmental stages that generally correspond to the early, middle and late stages of OFT remodeling and septation (47, 49 and 51 pairs of somites). The large-scale single-cell data empowered us to unbiasedly and systematically dissect the cellular diversity and heterogeneity during OFT development. We identified 17 cell clusters that could be assigned to six cell lineages. Among these lineages, the macrophage and VSMC lineages of the developing OFT have seldom been previously described in detail. In accordance with the myocardial to arterial phenotypic change, we observed the myocardial lineage diminished over time, while the VSMC lineage expanded during development. We provided molecular signatures for the cell lineages and clusters, and highlighted that *Cxcl12* could serve as a specific early marker for the embryonic VSMC lineage at the base of the great arteries. Cell lineage relationships and cellular transitions, such as EndoMT, have been identified through analyzing the dynamic changes in cell states by a force-directed layout of the KNN graph. In particular, we identified convergent development of the VSMCs at the base of the great arteries that has not been recognized before, where intermediate cell subpopulations were found to be involved in either myocardial to VSMC trans-differentiation or mesenchymal to VSMC transition. Through smFISH, we observed that cells expressing the myocardial marker *Myh7* co-expressed various levels of *Bmp4* (the marker gene for the myocardial c9 cluster) and the VSMC marker gene *Cxcl12* in OFT walls, thus confirming the existence of myocardial to VSMC trans-differentiation. Moreover, we found that the *Penk*^+^ cluster c8, a relatively small mesenchymal subpopulation that was undergoing mesenchymal to VSMC transition, was specifically associated with the fusion of the OFT cushions. Through pseudo-temporal ordering and GRN analysis, we uncovered the expression dynamics and critical transcriptional regulators potentially governing cell state transitions during OFT development. Finally, we developed web-based interactive interfaces for our dataset to facilitate further data exploration.

### Cellular diversity of developing cardiac OFT uncovered by large-scale single-cell profiling

Defining the lineage, proportion and molecular signature of distinct cell types is fundamental to our understanding of developmental processes^43^. Single-cell RNA-seq has revolutionized developmental biology by allowing for unbiased and systematic characterization of the cellular states in developing systems, such as the developing human fetal kidney^44^ and prefrontal cortex^45^. In a study on single-cell anatomical mapping of the embryonic heart^19^, the authors investigated the cellular composition and gene signatures of the OFT. However, only a total of 371 OFT cells (E10.5) were analyzed, which may be insufficient for a detailed dissection of heterogeneity. Apart from the four cell lineages (myocardial, epicardial, endocardial and mesenchymal) previously described^19^, our large-scale single-cell RNA-seq empowered us to detect a relatively rare (0.5%) cell lineage, i.e., macrophages (Figure 2B). Notably, the relative proportion of macrophages did not significantly change during development (Figure 2C), implying their important role in OFT development. Given that apoptosis is a ubiquitous process during development including OFT development^46^, macrophages are thought to function to remove debris arising from normal apoptosis. Nevertheless, it has been increasingly recognized that macrophages residing in tissues play essential roles in normal development. For example, macrophages are required for coronary development via mediating the remodeling of the primitive coronary plexus^47^. Our findings highlight the role of macrophages in OFT remodeling and suggest avenues for further investigation into the role of macrophages in cardiac development.

We also characterized the VSMC lineage of the developing OFT which has not been described in previous studies^19^. Our data showed that the VSMC lineage constituted the second largest (22.3%) cell population of the developing OFT (Figure 2B) and significantly expanded over development (Figure 2C). This observation is in line with our knowledge that OFT walls undergo myocardial to arterial phenotype change during development^13^. Although *Cxcl12* was previously found highly expressed in the walls of the aorta and pulmonary trunk of the embryonic heart (E12.5)^28^, our data highlights that *Cxcl12* could serve as a specific early marker for the embryonic VSMC lineage of the great arteries (Figure 2D). Through smFISH, the expression of *Cxcl12* was observed specifically in cells that would eventually form mature smooth muscle structures. For example, *Cxcl12* was highly expressed in a strip of cushion mesenchymal cells between the lumens of aorta and pulmonary artery, which would develop into the aorticopulmonary septum, a smooth muscle structure that eventually forms the facing walls of the great arteries (Figure 6A). The chemokine Cxcl12, which is secreted mainly in smooth muscle cells, has been suggested to be essential for coronary artery development through driving migration of cells expressing its receptor Cxcr4, e.g., endothelia cells^28^. Cxcl12-Cxcr4 signaling has also been suggested to be required for correct patterning of pulmonary and aortic arch arteries possibly by protecting arteries from uncontrolled sprouting^48^. Although the mechanism underlying the arterial system development mediated by Cxcl12-Cxcr4 signaling remains elusive, our result may suggest a role in septation and remodeling of the OFT.

### Intra-lineage heterogeneity of developing cardiac OFT unraveled by large-scale single-cell profiling

Cellular heterogeneity is a general feature of biological tissues and exists even within seemingly ‘homogeneous’ cell populations^16^. The large-scale single-cell RNA-seq unraveled previously unrecognized cellular heterogeneity within each cell lineage of the developing OFT. Except for macrophages, all other five cell lineages displayed distinct cell clusters/subpopulations (Figure 1B, Figure 2B). *Tmem255a,* previously reported to be an epicardial marker of the embryonic heart^19^, was found only mark one of the two subpopulations of the OFT epicardial lineage (Figure 2D, Online Figure IV). Samples derived from each developmental stage were found to have cells in almost all of these clusters, and they differ only in terms of relative proportions (Figure 1C). This result illustrates that each of our samples captured a full spectrum of cellular states throughout development due to cellular asynchrony. This finding also gave us a unique chance to identify the intermediate, transitioning subpopulations that have not been characterized before. For example, we identified c9 as a myocardial subpopulation undergoing myocardium to VSMC trans-differentiation (Figure 4). Additionally, the transcriptomic heterogeneity among subpopulations was predominately driven by the cellular positions along the transition, differentiation or maturation of cell lineages, as reflected by the dynamic changes in relative proportions over time for many clusters (Figure 1C, Figure 3C). For example, the VSMC cluster c3 rapidly expanded over development and represented a more mature state of the VSMC lineage than c4 did. Nevertheless, transcriptomic heterogeneity may also be influenced by other factors. For example, c8 represents a small mesenchymal subpopulation associated with the fusion of OFT cushions.

### Myocardial to VSMC trans-differentiation occurred during the OFT development

From the onset of development, the OFT is encased by a myocardial wall. As the progression of septation and remodeling, the myocardial wall rapidly changed into an arterial phenotype, characterized by the trick layer of smooth muscle cells in the tunica media. This myocardial to arterial phenotypic change has been previously described^13, 49^, and our smFISH with probes marking the myocardial (*Myh7*) and VSMC (*Cxcl12*) lineages on serial sections of the OFT clearly displayed this process (Figure 6A). However, the fate of the myocardium during this process remains controversial to date. It has been suggested that trans-differentiation of myocardial cells to arterial components may occur during OFT development in the embryonic hearts of chicken^50^, rat^49^ and mouse^51^. Conversely, it is also hold that the phenotypic change could just be considered a regression of the myocardium^13^. Our single-cell dataset supports the view of myocardial to VSMC trans-differentiation by identifying cell clusters representing a continuum of cell state transitions (Figure 3D). Expression profile comparison analysis demonstrated that myocardial cluster c9 and VSMC cluster c4 were in an intermediate state along the trajectory of myocardium to VSMC trans-differentiation (Figure 4). Through smFISH we observed that cells expressing myocardial marker *Myh7* co-expressed various levels of *Bmp4* (c9 marker gene) and VSMC marker gene *Cxcl12* in OFT walls, thus confirming the myocardial to VSMC trans-differentiation (Figure 6). A recent study demonstrated that myocardial cells may transit to the mesenchymal cells of the intercalated cushions during OFT development^52^. Thus, our findings provide additional evidence that highlights the plasticity of the embryonic myocardial cells of the OFT. All these findings imply that transitions between cell lineages during OFT development may be more complicated than previously appreciated.

### Convergent development of the VSMCs at the base of the great arteries

Cell lineage relationships and cellular state transitions can be inferred from time-series single-cell transcriptomic data even for a complex developmental system, such as embryogenesis of frog^53^ and zebrafish^54^, through a force-directed layout of the KNN graph. Based on the dynamic change in cell states over time reflected by the KNN graph of our time-series single-cell data, known cell transition events were recapitulated, for example, EndoMT that underwent between the endocardial and mesenchymal subpopulations (Online Figure VIA). In particular, we identified convergent development of the VSMCs at the base of the great arteries that has not been recognized before, where intermediate cell subpopulations were found to be involved in either myocardial to VSMC trans-differentiation or mesenchymal to VSMC transition (Figure 3F & 3G). We found that the mesenchymal to VSMC transition involved one mesenchymal subpopulation, c1, that occurred mainly at the late stage, and another smaller mesenchymal subpopulation, c8, that occurred mainly at the early stage. Such a temporal relationship for mesenchymal to VSMC transition has never been recognized before. Furthermore, by smFISH, the c8 *Penk*^+^ subpopulation was found to be specifically associated with the fusion of OFT cushions, in line with its relatively small size and populating the early stage samples (Figure 6C). In addition to the mesenchymal subpopulations, the myocardial subpopulation c9 also contributed to the development of the VSMC lineage. Altogether, three developmental paths were identified to be implicated in convergent development of VSMC lineage at the base of the great arteries, which involves different cell lineages and different subpopulations of the same lineage.

Furthermore, by pseudo-temporal ordering and GRN analysis, we uncovered gene expression dynamics and critical transcriptional regulators potentially governing the cell state transitions during the development of VSMCs (Figure 5, Online Figure V). The Notch signaling pathway has been known to positively regulate the specification, differentiation, and maturation of VSMCs^38^. We found that the expression of genes in the Notch signaling pathway, including receptor (*Notch1*), ligand (*Jag1*) and downstream targets (*Hey1*, *Hey2*, *Heyl* and *Pdgfrb*), was positively correlated with the reconstructed trajectories for all the three paths. Then, we identified *Heyl*, encoding a known downstream TF of the Notch signaling, at the top of the upregulated regulators. Thus, our results highlight the role of the Notch signaling pathway in the development of the OFT VSMC lineage. We provide a set of critical transcriptional regulators that were sequentially activated or repressed along the development trajectory. For many of these regulators, the roles in VSMC development have seldom been suggested before.

In conclusion, through large-scale single-cell transcriptomic sequencing, we performed an unbiased and systematic study on the cellular types and states of the cardiac OFT during development. Our results support the existence of myocardial to VSMC trans-differentiation, and convergent development of the VSMC lineage at the base of the great arteries. We provide a single-cell reference map of cell states for normal OFT development, which allows the CHD community to assess how perturbations affect the transcriptomic states of OFT cell lineages, to prioritize candidate genes of OFT malformations based on expression, and to map candidate genes to cell types or subpopulations. Our study demonstrated the power of time-series single-cell transcriptomic data for identifying cell state transitions in a complex developmental system.

## ACKNOWLEDGMENTS

We thank the staff in the public platform of the State Key Laboratory of Cardiovascular Disease for providing technical assistance.

## Nonstandard Abbreviations and Acronyms

EndoMT: endothelial to mesenchymal transition
GRN: gene regulatory network
KNN: k nearest neighbor
OFT: outflow tract
SHF: second heart field
UMI: unique molecular identifier
VSMC: vascular smooth muscle cell

